# Controlling the Formation of Multiple Condensates in the Synapse

**DOI:** 10.1101/2025.02.19.638975

**Authors:** Nahid Safari, Maria Augusta do Rego Barros Fernandes Lima, Riccardo Rossetto, David Zwicker, Elisa D’Este, Christian Tetzlaff

**Author notes:** Contributing authors.

## Abstract

The postsynaptic density (PSD) is a molecule rich structure that continuously adapts its organization controlling synaptic physiology and transmission. Experimental studies have shown that this organization is dominated by the formation of condensates or (nano)clusters and by the seamless transition in their numbers as response to synaptic plasticity. In this study, we utilize different computational modeling frameworks to show that variations in the level of local protein concentrations together with the modulation of protein binding strengths can be key molecular factors in controlling the formation and number of clusters in the synapse. Comparison to MINFLUX data of spatial localization of PSD95 in the postsynapse under different activity conditions allowed us to derive predictions about the correlation between these two factors across a population of synapses. Co-variations of the factors, to mimic a simple LTP protocol, shows that a PSD containing a single cluster can reorganize to form multiple clusters that persists for long periods of time, providing a potential explanation of recent experimental data of PSD reorganization.

## Introduction

The postsynaptic density (PSD) is a key membraneless organelle of the neuronal synapse essential for signal transmission and memory formation. It contains hundreds of different proteins, among which scaffolding proteins, receptors, actin filaments, and adhesion molecules [1, 2, 3]. Although there are detailed insights into individual PSD components, the dynamic regulation of these components, particularly the positioning of AMPARs and scaffolding proteins like PSD95, remain unclear [4, 5]. Recent studies suggest that the organization of the PSD is mainly governed by liquid-liquid phase separation (LLPS) of PSD-related molecules, forming condensates or dense molecule clusters within a dilute phase.

Indeed, some *in vitro* reconstitution experiments have demonstrated that the mixing of primary scaffolding proteins can result in the formation of molecule clusters through LLPS, suggesting that PSDs are dynamic structures where components are in constant flux [6, 7, 8]. Several factors can influence the formation of LLPS-induced molecule clusters, as the concentration of client molecules or the presence and number of intrinsically disordered regions (IDRs). External factors such as temperature, pH-level, or salt concentration, also affect the propensity of molecules to form clusters [9, 6, 4]. Despite the progresses being made in understanding these processes, the interplay between different internal and external factors in determining LLPS and the formation of (multi)clusters still requires detailed investigations.

In cells, PSD as visualized by fluorescent labeling of their key components (e.g. PSD95) appear as clusters of few hundreds of nanometers in size. Within minutes after the application of chemical long-term potentiation (cLTP) protocols, the PSD reorganizes in becoming more complex, with perforation and segmentation of single clusters into multiple clusters [10, 11, 12]. This initial phase is followed by the accumulation of scaffolding proteins (e.g. PSD95) and AMPA receptor recruitment [13, 14], a process critical for maintaining synaptic changes and supported by new protein synthesis [15, 16]. However, the mechanisms driving PSD reorganization and the impact of new protein synthesis on PSD structure remain unresolved.

Standard mathematical models of cluster formation typically exhibit two outcomes; either no clusters form or multiple clusters eventually merge into a single cluster, via Ostwald ripening or Brownian coalescence [17]. Brownian coalescence relies on cluster motion, which is suppressed through membrane anchoring in PSDs. In contrast, Ostwald ripening can be prevented by continuous dissociation of cluster-forming molecules in combination with a steady influx of new proteins. Studies using concentration field models of droplet formation [18, 19] or reaction-diffusion models of inhibitory synapses [20, 21, 22] showed that both processes can be in balance such that the system always contains multiclusters. Similar to these ideas, by combining computational modeling with experimental data, in this study we reveal that the fine-tuned interplay between intracellular concentrations of PSD-related molecules and their inter-molecular binding strengths can lead to the formation of multiple PSD-clusters that are maintained for a sufficient duration. As these two factors can be regulated by cellular processes like local protein synthesis or co-factor concentrations, a neuron could use them to “control” the number of PSD clusters being formed in a synapse for proper synaptic functioning.

## Results

### Formation of multicluster postsynaptic density

To elucidate the mechanisms underlying PSD organization, we consider a simulation box of about 0.001*µm*^3^ (1% to 10% of average spine volume) that resembles the PSD within a dendritic spine. As minimal mechanistic model of protein dynamics, we focus on the spatio-temporal dynamics of the scaffolding protein PSD95, being one of the most prevalent protein in the PSD. As balance between computational efficiency and molecular realism, similar to previous modeling approaches [23], we utilized a minimal coarse-grained model of a PSD95 protein, meaning that it is modelled as a Brownian particle consisting of a core structure with a diameter of 5*nm* [24, 25] and fixed number of molecular interaction sites or patches that represent PSD95’s binding domains. To reduce the complexity of the partially unknown network of molecular interactions that directly and indirectly interlinks several PSD95 proteins, we integrate these interactions into direct particle-particle bindings, and validate the model by matching computational results [23] of the phase space of LLPS-dependent cluster formation given the periodic boundary condition (see supporting information).

To enable analyses of the influence of processes like protein synthesis on clustering dynamics, we adjusted the boundary condition to the so-called fixed-concentration boundary condition. Fixed-concentration boundaries imply, different to the often used periodic boundary condition, that the simulation box is being linked to a “virtual environment” [26]. Thus, by crossing the boundary particles can move out of the simulation box, while new particles can enter the simulation box depending on the preset particle concentration *C*_*env*_ in the virtual environment. As the resulting in- and out-flux of particles at a boundary segment depends on the dynamics of particles in the immediate vicinity of the segment, even after a long equilibration time, complex particle flows can result in a continuous exchange of particles between simulation box and environment. Within the environment, we do not simulate individual particles. Note that we only consider the immediate vicinity of the PSD as virtual environment (Figure 1A), assuming that all other neuronal compartments influence the PSD through changes in the virtual environment. The latter implies that protein synthesis, occurring at various sites like the cell body, dendritic shaft, or spine head near the PSD, leads in the model to variations in particle concentration of the virtual environment. Therefore, our system recapitulates experimental evidences according to which protein synthesis is a tool utilized by neurons to tune synaptic plasticity.

**Fig. 1:**
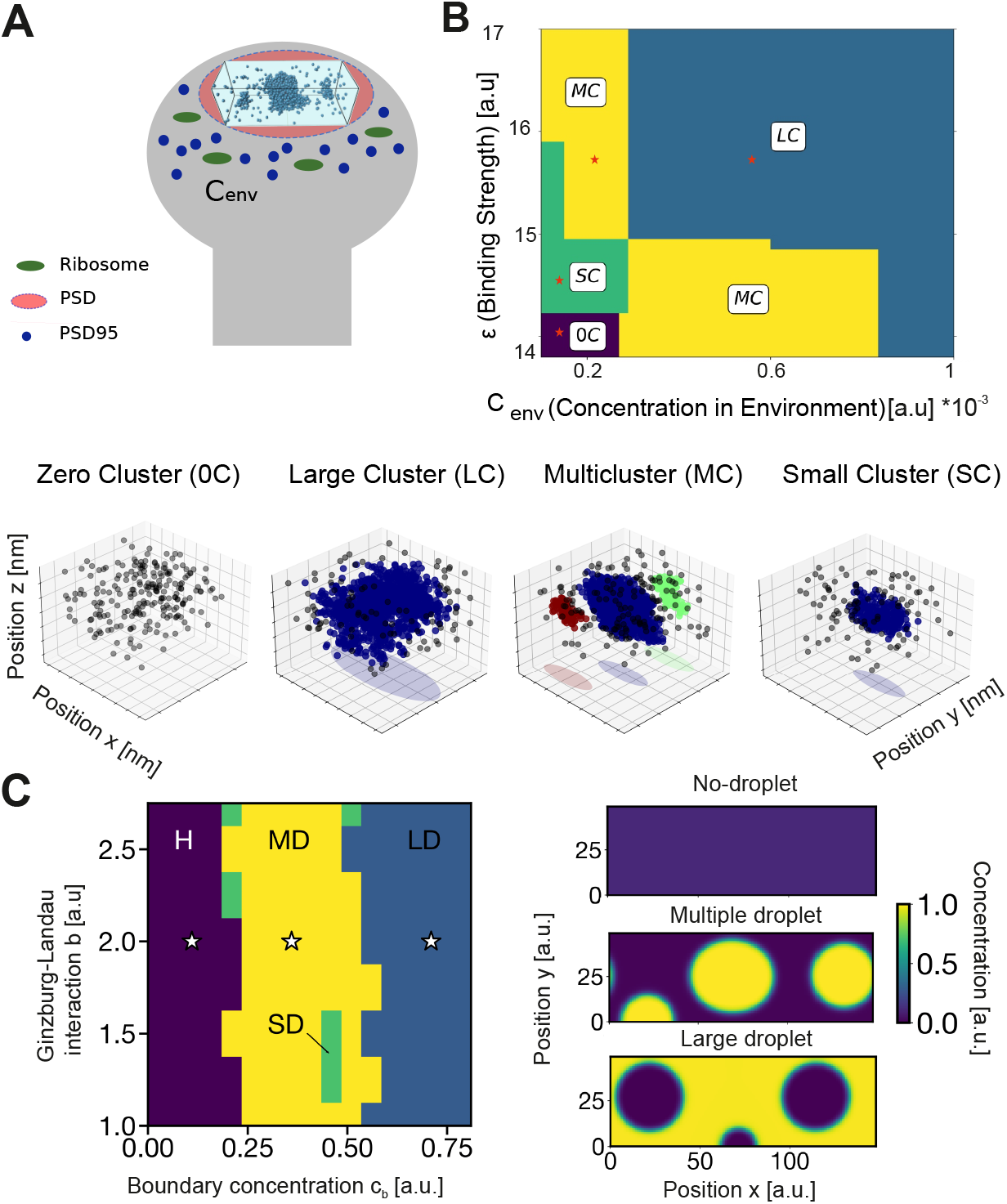
Schematic of computational model and resulting phase diagram. A) In the particle-based simulation of PSD organization, we simulate dynamics within a box of 0.001*µm*^3^ that resembles the intracellular volume PSD. Proteins, being synthesized by ribosomes, influence the concentration of particles *C*_*env*_ outside the simulation box. Depending on *C*_*env*_, new particles can enter the simulation, while particle that hit the simulation box boundary, leave the simulation. B) The phase diagram of the particle-based simulation in dependence on the particle concentration in the immediate environment *C*_*env*_ and particle-particle binding strength *ϵ* shows the emergence of different phases: zero cluster (0C) phase (purple), small cluster (SC) phase (green), large cluster (LC) phase (blue), and multicluster (MC) phase (yellow). Pictures on the right-hand side show snapshots of the particle organization in different phases, with clusters detected by DBSCAN in different colors. C) The phase diagram of a field-theoretic model with boundary reactions shows the emergence of a no-droplet phase, large droplet phase, and multiple droplet phase. The phase diagram is obtained evolving the dynamics from a randomly perturbed homogeneous state with concentration *c* = *c*_0_, and stopped after 10^4^ time steps. Remaining parameters are *L* = 37*ℓ*_0_ and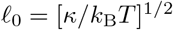, where 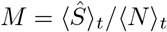and *t*_0_ = *κ/*(*k*_B_*T*)^2^*M*. See Methods for details.

Simulations are executed following standard procedures from molecular dynamics simulations: After equilibrating the system with periodic boundary condition into a liquid phase, the simulation box is elongated, and simulations at specific binding strengths reveal liquid-liquid phase separation (LLPS). Using the LLPS state as initial condition, the boundary condition is switched from the periodic condition to the fixed concentration condition with given concentration *C*_*env*_ and binding strength *ϵ*. This process is repeated for different *Cenv* and*ϵ*, revealing the states shown in Figure 1B:

#### Zero Cluster (0C) state

For low values of both *ϵ* and *C*_*env*_, the limited number of available particles prevents them to interact sufficiently to form a cluster. *Large Cluster (LC) state*: For high values of *ϵ* and *C*_*env*_, the majority of proteins aggregate into a large cluster that occupies most of the simulation box, leaving only a small number of particles in the dilute phase.

#### Small Cluster (SC) state

For low *C*_*env*_ and intermediate values of *ϵ*, we observe a different type of single cluster emerging in the system, which differs from the large cluster being formed in the LC state. Here, due to the lower binding strength only a small number of particles aggregate into the cluster, while the majority of particles remain as free particles in the surrounding dilute phase.

#### Multicluster (MC) state

For certain value regimes of *ϵ* and *C*_*env*_ (yellow areas in Figure 1B), multiple clusters are formed by the particles and remain stable for long time intervals. Given the intrinsic stochasticity of the system, the number of clusters can vary on a slow timescale (see supporting information).

To verify our mathematically complex particle-based simulations with nonstandard boundary condition, we employed a field-theoretic model for droplet formation. The model is based on Cahn-Hilliard dynamics with an underlying Ginzburg-Landau free energy and incorporates boundary reactions to mimic the fixed-concentration boundary condition. Similar to the particle-based simulations, the phase diagram of the field-theoretic model shows the emergence of a pure dilute phase, a large droplet phase, and a phase with multiple droplets as a function of the interaction parameter resembling the particle-particle interaction, and the boundary concentration akin to the particle concentration in the environment (Figure 1C). Differences between the resulting phase diagrams can be expected given that parameters between models cannot be matched in a one-to-one fashion.

Investigating the differences between states in more detail, first, we surveyed the temporal development of the system. In contrast to simulations with periodic boundary condition, which implies a constant number of particles, the fixed-concentration boundary condition allows a continuous exchange of particles with the environment. Despite the continuous flow of particles, the number of particles reaches a stable mean number of particles ⟨*N*⟩ _*t*_ for the different states with ongoing minor fluctuations (Figure 2A, average across a time window of length *0*.*5 ns*). Although the examples shown in Figure 2A imply a relation between number of particles in quasi-static state, the number of particles does not conclusively predict the state of the system and, thus, the number of clusters (Figure 2B).

**Fig. 2:**
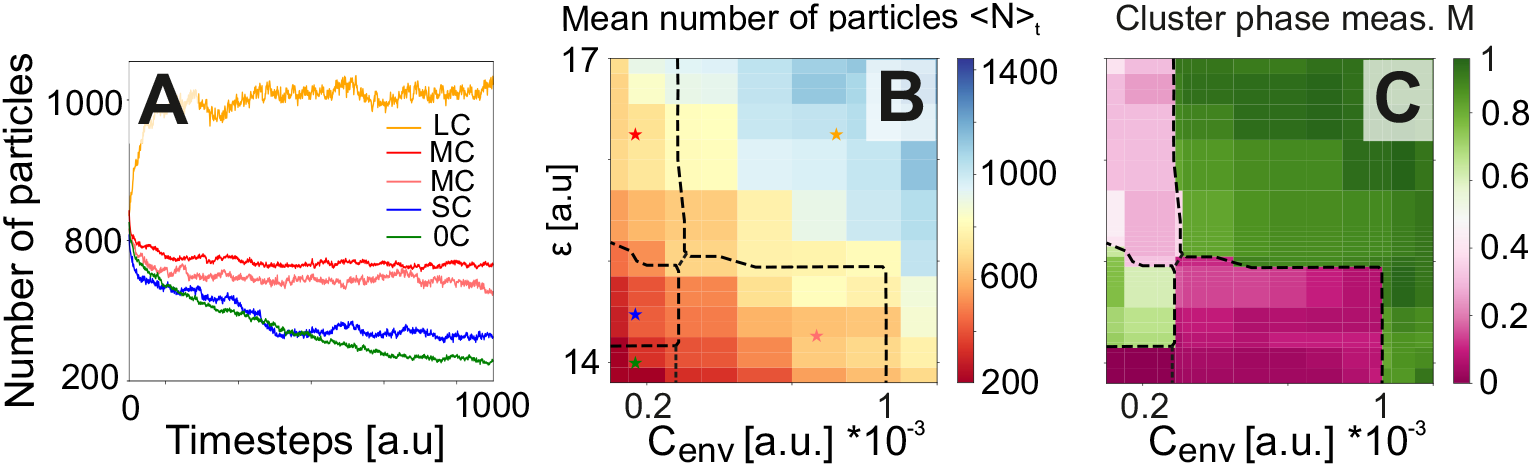
Evaluation of different states in particle-based simulations. A) Examples of the temporal development of the number of particles being in the simulation box for different values of *ϵ* and *C*_*env*_. Values are depicted in panel B). B) The mean number of particles being in the simulation box after relaxation. Borders of different states from Figure 1B are indicated as black dashed lines. C) The cluster state measure *M* for different values of *ϵ* and *C*_*env*_, after relaxation of the number of particles.

Next, we considered the average number of particles across all formed clusters *n*_*c*_ in the same time interval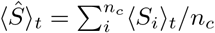, divided by the total number of particles

⟨*N*⟩ _*t*_, yielding the cluster state measure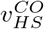. *M* is a number between zero and one, with *M* = 0 if no cluster is being formed, and *M* = 1 if one cluster contains all particles being in the simulation box. Calculating *M* for all cases of different values of particle concentration in the environment *C*_*env*_ and particle-particle binding strength *ϵ* shows that the different states can be clearly identified (Figure 2C).

### Model-based prediction of activity-dependent correlations between external and internal factors controlling cluster number in synapses

Our results indicate that the cluster states depends on the particle concentration in the environment *C*_*env*_ and on the particle-particle binding strength *ϵ*. To test whether there is a dependency between these two variables, we analysed experimental data of synapses under different activity conditions. In particular, we utilized a dataset rendering PSD95 in dendritic spines with 3D MINFLUX, a nanoscopy technique capable of localizing molecules with 5nm isotropic resolution (i.e. the size of our simulated particles) [27]. Thereby, the dataset provides information regarding the distribution of the PSD95 molecules both at the PSD itself and in the space in proximity, which resembles our virtual box. Lastly, the dataset contains information recorded from hippocampal primary neurons in control condition or treated with tetrodotoxin (TTX) to block neuronal activity, using gabazine (GZ) to enhance neuronal activity for 24 h. In this timeframe, both PSD structure and protein synthesis, our external factor influencing *C*_*env*_, are modulated to react to the activity change.

Similar to the analyses of particle-based simulation data, we applied the DBSCAN clustering algorithm to identify PSD clusters within individual spines of the experimental data (see supporting information for analysis details). As expected from literature and simulations, we identified dendritic spines hosting PSDs being in the SC state, LC state, and MC state (see examples in Figure 3A). A subsequent calculation of the cluster state measure *M* reveals that in the control condition about 61.0%± 0.8% of dendritic spines have multiclustered PSDs, while 17.1%± 2.1% have large, single clusters and 21.9%± 2.5% have small, single cluster (Figure 3Bi). To compare the experimental findings with our simulations, we consider that each synapse has a slightly different molecular composition [28] implying different values for *ϵ* and *C*_*env*_. Therefore, we draw for each model spine values for *ϵ* and *C*_*env*_ from a Bivariate Normal Distribution (BND), with means *µ*_*ϵ*_, *µ*_*c*_, variances *σ*_*ϵ*_, *σ*_*c*_, and correlation coefficient *ρ*. We utilized the optimization framework Optuna [29] to find the values of the BND-parameters such that the resulting distribution of cluster states for the model synapses (blue dots in Figure 3C; neglecting the 0C state) best match the frequency distribution of cluster states from experiments (see supplementary information for more details). The resulting value of *ρ* (indicated by the red line in Figure 3C) provides the prediction of the expected correlation between external factor *C*_*env*_ and internal factor *ϵ* across all synapses in the experimental data under given condition. For the control condition, this results to *ρ* = −0.29, meaning a weak negative correlation or, in other words, in a single synapse, if *C*_*env*_ is high, then *ϵ* tends to be low, and vice versa.

**Fig. 3:**
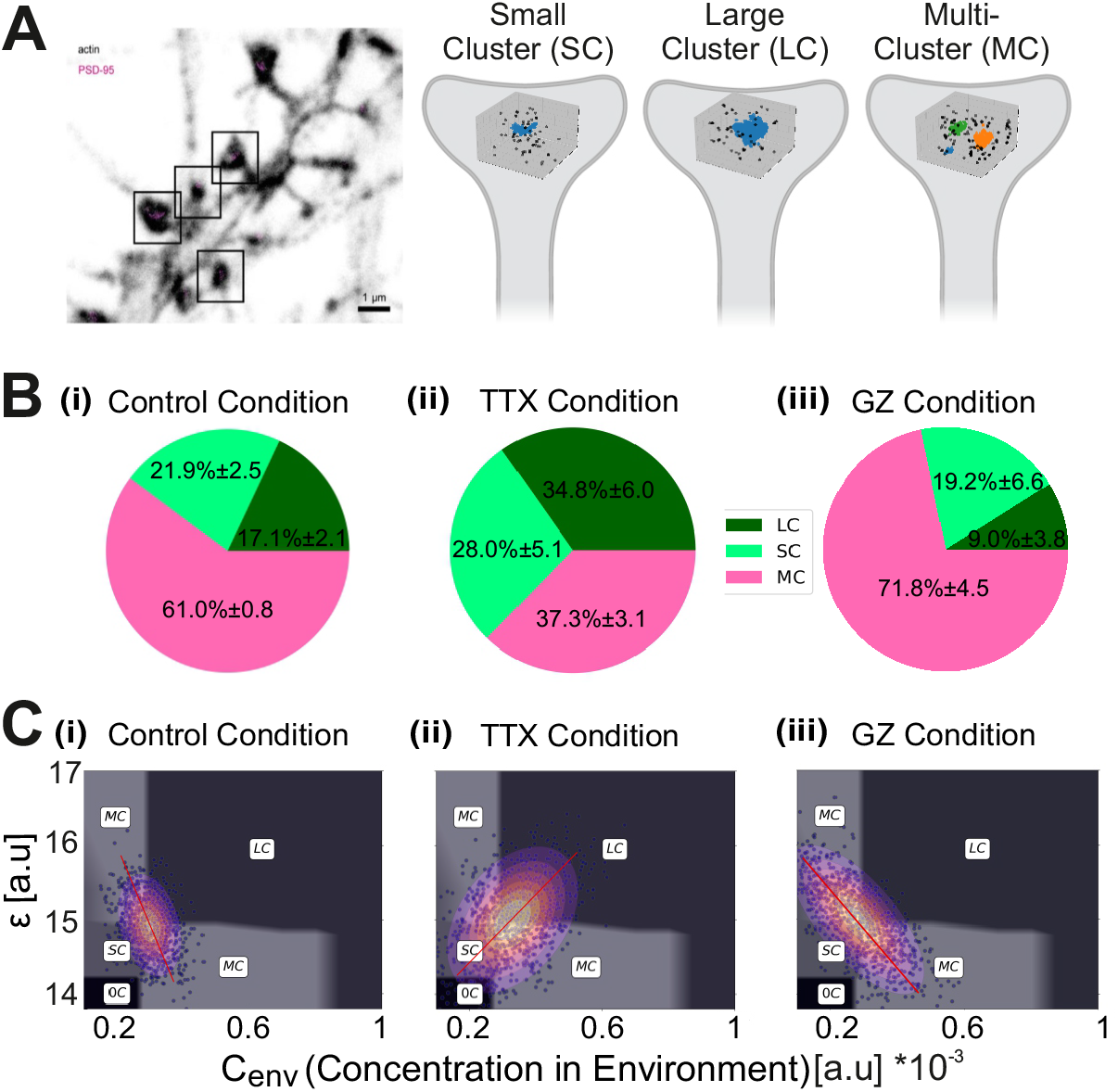
Prediction of correlation between *ϵ* and *C*_*env*_ by comparison to 3D MINFLUX data. A) The experimental data shows the spatial distribution of PSD95, labeled with a nanobody in magenta (DNA-PAINT 3D MINFLUX), in dendritic spines. Using DBSCAN clustering on PSD95 protein coordinates, we identified distinct states similar to particle-based simulations. B) The distribution of dendritic spines in different cluster states depends on the treatment condition: (i) control, (ii) TTX, (iii) GZ. C) Matching the simulation results with the experimental data shown in panel B leads to the predicted correlation (red line) between *ϵ* and *C*_*env*_. In gray is shown the phase diagram from Figure 1B. Blue dots indicate example model synapses with parameters drawn from the fitted BND, which is shown as contour graph. (i) Control: *ρ* = −0.29; (ii) TTX: *ρ* = 0.40; (iii) GZ: *ρ* = −0.80.

Prolonged exposure to TTX causes PSD changes aiming at maximizing the efficiency of signal trasmission (homeostatic plasticity). When analysing data derived from TTX-treated neurons, we observe that the number of PSDs in the multicluster state reduces significantly to 37.3%± 3.1%, while the number of spines in the large cluster state increases to 34.8%± 6.0% (Figure 3Bii), in line with previous observations. Comparison to the particle-based simulation reveals that these changes could result from a larger variance in values of *C*_*env*_ and *ϵ* and a positive correlation between both factors (*ρ* = 0.40). The latter implies that a PSD having more proteins in its adjacent environment, also experiences a higher level of protein-protein interaction [30, 31] (Figure 3Cii).

Interestingly, GZ-induced activity enhancement cause the opposite effect on PSDs, with the number of PSDs in the multicluster state increases to 71.8%± 4.5% compared to control conditions, while the number of single, large clusters is reduced to 9.0% ± 3.8% (Figure 3Biii). Under this condition, the comparison to the simulation results shows that the correlation coefficient between *C*_*env*_ and *ϵ* is strongly negative (*ρ* = −0.80; Figure 3Ciii). Thus, our analysis yields the prediction that increased activity levels leads to an inverse correlation, meaning that dendritic spines with a high number of proteins in the environment of the PSD have a low level of protein-protein interaction and vice versa. Overall, these data show that our model is able to accurately predict molecular changes (*ϵ*) that are invisible when analyzing the morphology of PSDs alone, and reveal a strong activity-dependent modulation of parameters.

### Controlled transitions between cluster states

The induction of long-term synaptic potentiation (LTP) triggers a cascade of diverse molecular processes, distributed across a wide variety of timescales [32, 14, 33, 34, 35]. Immediately after LTP induction, there is documented evidence of changes in PSD morphology [12], likely happening without an increase in protein synthesis rate [36, 37, 38]. About one hour after LTP induction, a noticeable increase in the abundance of PSD95 proteins is being reported [13, 14, 39].

Inspired by these findings, we implemented a minimal model of transient molecular changes, adapting *ϵ* and *C*_*env*_ in a cascade like manner (Figure 4A). First, we initialized the system with parameter values that lead to a single, small cluster state (state ‘0’, Figure 4B), representing the basic state. Then, we trigger a cascade by first increasing the particle-particle interaction *ϵ* (state ‘1’). As expected from the phase diagram (Figure 1B), the system reaches the MC state, forming several cluster (indicated by different colors in Figure 4B). Considering the delayed increase in protein synthesis rate, afterwards we also increase the number of particles being in the environment *C*_*env*_ (state ‘2’), leading to a transition to the LC state. As most molecular processes after LTP induction are transient, we assume that *ϵ* goes back to its initial value, while an increased *C*_*env*_ value persists (state ‘3’). This change in parameters leads to a state transition to the MC state, as expected from the system’s phase diagram. Last, also *C*_*env*_ goes back to the basic state value, such that parameters reach state ‘0’ again. Although parameters have the same values as in the basic state, after the complete cascade of transitions, the system stays in the MC state, with significant difference in *M* and number of clusters across several trials (Figure 4C). Thus, this cascade introduces a lasting change in cluster state, moving from single to multiple clusters, like LTP results to the emergence of more multiclustered PSDs [11, 12].

**Fig. 4:**
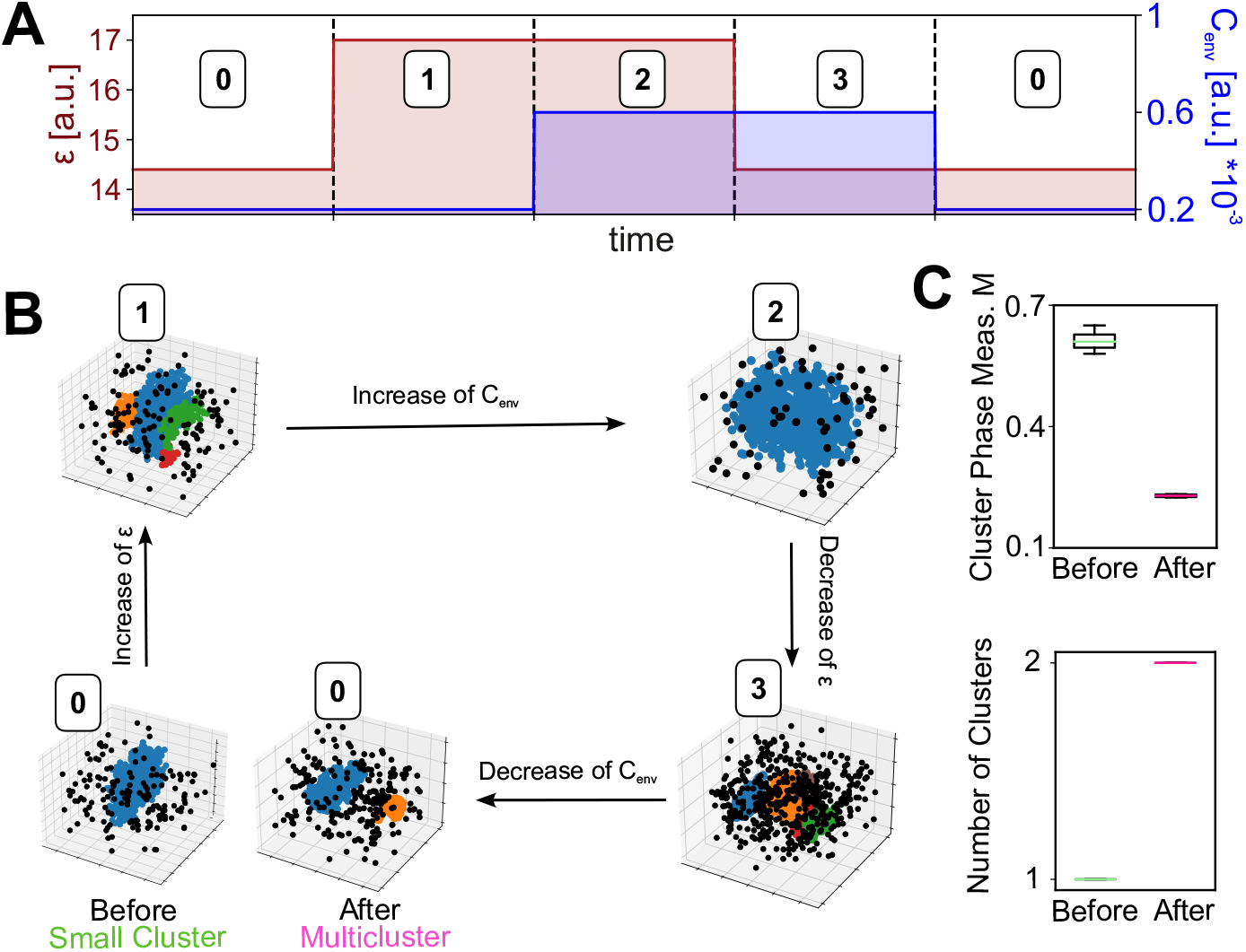
Transient changes of *ϵ* and *C*_*env*_ trigger lasting state transitions. A) Protocol of cascade-like changes in values of *ϵ* and *C*_*env*_. Note that for each pair of values, we let the particle-based simulation relay into its new state. B) One example of state transitions given the protocol shown in panel A). C) Average value of cluster state measure *M* and number of clusters *NC* for five separate trials, before applying the cascade of state transitions (‘before’) and after completing the whole protocol (‘after’).

## Discussion

In this study we integrated different computational model frameworks with experimental data to investigate different factors that determine the clustering of molecules in membraneless compartments like the postsynaptic density. Several studies using patchy particle models have highlighted the crucial role of protein-protein interactions [23, 40, 41, 42] in organizing membraneless compartments, yet many overlook the fundamental influence of external factors changing protein concentration. Our research addresses this gap by showing how variations in external particle concentrations in the immediate vicinity of the area of interest influence the formation of distinct organizational clustering states. While protein-protein interactions are essential, their interplay with environmental conditions can lead to diverse patterns, including the formation of a multicluster state with multiple coexisting clusters, as measured in the PSD [43, 10, 44, 45]. For this, in the model, the continuous exchange of particles with the environment is essential for the emergence of the multicluster state as well as for its maintenance, as changing the boundary condition after multicluster formation to a periodic condition leads to a perturbation of the state (see Supporting Information). Furthermore, over a longer timescale, multiple clusters may merge (see Supporting Information). The complex, continuous exchange of particles imposed by the fixed-concentration boundary condition, impedes the derivation of mathematical solutions to support the numerically found system’s phase diagram, including the emergence of the MC state and its stability. However, our numerical results are supported by a field-theoretic model with similar boundary condition. This model provides a promising candidate for further mathematical analyses of the system’s phase diagram to obtain a better understanding of the influences of different internal and external factors, and of their interplay.

The consideration of a fixed-concentration boundary condition is motivated by biological processes in neurons. Indeed, neurons can fine-tune protein concentration in several ways. Within minutes after activity stimulation, the volume of dendritic spines increases and the protein degradation machinery can be activated, both resulting in an effective lower protein concentration. New protein synthesis has the opposite effect and happens as quickly as 1.5 hours [46]. Therefore, by incorporating a fixed-concentration boundary condition and allowing proteins to move between the simulated compartment and the surrounding environment, our model reflects the physiological dynamic regulation in protein concentration. [36, 47].

In our study, we conducted molecular dynamics simulations using particles characterized by low viscosity to replicate the dynamics of particle movement in aqueous environments. By simulating the particles in this manner, we gain insights into the interactions and behaviors that occur when PSD95 proteins are suspended in a liquid medium. Of course, the dendritic spine and the PSD are much more complex molecular structures, including a large variety of protein interactions, modifications, and local differences in concentrations [28]. Here, we simplify this complex network by considering only one particle type with particle-particle interactions implemented by specific patches. Changes of the number of patches changes the parameter values for state transition, but maintains the overall structure of the phase diagram (see supplementary information). Further analyses are required to understand the influence of the different mixtures of proteins, for which the present model can be extended similar to a previous study of condensate formation with protein mixtures [23]. However, a more detail model of the clustering processes in the PSD requires a careful evaluation of parameters, likely asking for more experimental data.

By comparison to MINFLUX data, we predict an activity-dependence of the correlation between the factors of particle-particle interaction and environmental particle concentration. Note that the MINFLUX imaging technique might capture 50 % - 80 % of the actual protein population of all PSD95 molecules [27]. While the comparison between simulation and experiment are based on the visualized proteins, the subsampling of proteins is likely independent of protein density, such that we do not expect major discrepancies in the predicted correlations. Of course these activity-dependent correlations across population of synapses have to be tested, which is a demanding task, given the need to parallel measure number of proteins in the vicinity of the PSD and accessing the average protein-protein interaction. While the former can be done with state-of-the-art super-resolution fluorescence methods like MINFLUX [48] or ONE [49], the latter often requires the isolation of the protein of interest [50, 51], which is difficult in protein-rich areas like the PSD under different activity conditions. However, recent advancements in artificial intelligence show promising results [52], such that an interdisciplinary study could verify our predictions.

The long-term storage of information using a variable substrate like proteins is a difficult problem. Computational models suggest that this problem can be resolved by the collective dynamics of the involved proteins providing bistable or hysteretic system characteristics [53, 54, 55, 56, 57, 58]. Hereby, a system’s parameter is being changed, for instance as a result of LTP-induction, triggering a state transition. If the parameter falls back to its initial value, the system stays in the new state, memorizing the transition or LTP-event. Similar to this approach, our particle-based simulation shows that by varying the factors of particle-particle interaction and their environmental concentration in a sequential manner, the system can undergo several state transitions, ending in a multicluster state that persists despite the factors having again their initial values. Compared to the models discussed above the main difference is that our model requires a specific sequence of changes of two parameters. This could form a “write protection” mechanism against noisy variations of the parameters themselves, which are also implemented by noisy molecular processes. To test the implication of our findings on the storage of information, the different clustering states and their transitions can be transformed to a Markov chain. This can be integrated into abstract neuronal network models like the cascade model [33, 32], to investigate the functional implication of molecular clustering on cognitive processes like memory.

## Methods

### Particle-based model

Using the PyRID simulation framework [26], we considered a simulation box of size 222×74×74*nm* or about 0.001*µm*^3^ to model the cellular area of cluster formation. Particles are modeled as nearly hard spheres with a continuous attractive square-well interaction for defined patches (*λ*_*r*_ = 50, *λ*_*a*_ = 49). We used the pseudohard-sphere interaction (PHS)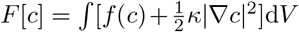 [59, 40] to describe the hard sphere behavior, ensuring that each particle occupies a given volume in the simulation box.

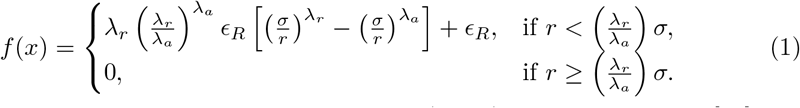

Furthermore, a continuous attractive square-well (CSW) interaction *v*_*CSW*_ [60] was applied to the associative sites or patches:

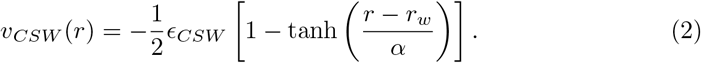

It is important to highlight that when referring to binding strength in the phase diagram, it specifically addresses *ϵ*_*CSW*_.

An important characteristic of such simulations is the considered boundary condition. The *periodic boundary condition (PBC)* maintains a constant number of particles throughout the simulation. PBC allows particles near the simulation box boundaries to exit from one edge and re-enter from the opposite edge. To account for the influence of the environmental concentration surrounding the simulation box (as in a biological cell), we considered the *fixed-concentration boundary condition*, allowing particles to move between the system and the environmental reservoir of particles.

Our model closely resembles previous studies of patchy particle simulations [40, 61]. To assess the phase diagrams, we employed two methods, the Direct Coexistence (DC) method [62, 63] and the molecular dynamics (MD) method [42, 64, 65]. The DC approach builds regularly extended slabs to represent the two phases — the condensed liquid (protein-enriched phase) and the diluted liquid (protein-depleted phase) — within the same simulation box. Using an NPT-ensemble MD method, equilibrium in the condensed-liquid phase was achieved. To ensure minimal vapor pressure, a high *ϵ*_*CSW*_ value (or low temperature) and a decreased pressure were maintained. The simulation box was then stretched along one axis to create the long side of the box (74· 3 = 222*nm*) after this equilibration. In order to investigate the phase behavior, several simulations were run in a constant NVT-ensemble, with the temperatures or the inverse protein-protein interaction strength (*ϵ*_*CSW*_) varied. Following a previous study [40], we set *α* = 0.01*σ* and *r*_*w*_ = 0.12*σ*. Due to the small size of *r*_*w*_, each individual patch has a valency of one, meaning each attractive site can only interact with one other patch at any given time. We set *σ* = 5*nm* being in the range of the measured size of PSD95 proteins. Once the LLPS phase is established, the boundary condition is adjusted from periodic to a fixed-concentration boundary condition.

### Fixed-concentration boundary condition

When studying the spatiotemporal organization of PSDs, it is essential to consider that protein synthesis in neurons is affected by neuronal activity, which persists over time. To capture the effects of this transient concentration change, we adjusted the boundary condition from periodic to fixed-concentration.

By implementing fixed-concentration boundary conditions, the simulation box is connected to an external reservoir of particles, allowing us to focus on a specific region of a larger system. This approach eliminates the need to simulate the dynamics of molecules outside the simulation box. In this setup, molecules outside the box are treated as “virtual,” entering the simulation only when they cross the boundary. The fixed concentration boundary condition in our model links the simulation box to a particle reservoir, enabling the simulation of the box within a larger system without directly modeling the dynamics of the external molecules. Virtual molecules only contribute to the simulation when they cross the boundary of the simulation box. The expected number of interactions (*N*_hit_) between a molecule type and the simulation box boundaries is calculated at each iteration, based on the fixed external concentration *C*, the diffusion coefficient *D*, and the total boundary surface area *A*. The average number of molecules that hit the boundary of area *A* from one side within a time step Δ*t* is computed using the molecule concentration *C* and the average distance a diffusing molecule travels normal to the plane, *l*_*n*_, within Δ*t* [66].

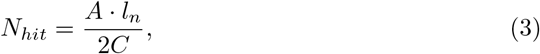

Where

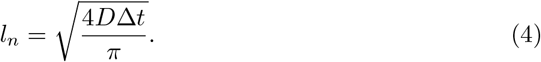

Here *D* = *Tr*(*D*^*tt*^)*/*3 is the scalar translational diffusion coefficient. As such, the number of molecules that cross the boundary each time step is drawn from a Poisson distribution with a rate *N*_*hit*_.

### Field-theoretic model

The field-theoretic model describes the evolution of the concentration field *c*(***r***, *t*) based on thermodynamic principles [67]. Assuming that particle count is conserved locally, we find *∂*_*t*_*c* = *M*∇ ^2^*µ* + *s*, where *M* represents the mobility coefficient, *s* denotes potential source/sink terms, and *µ* = *δF/δc* is the chemical potential of the described species, which follows from a suitable free energy *F*. We consider a two-dimensional, rectangular system with side lengths *L*_*x*_ and *L*_*y*_ along the respective coordinates *x* and *y*. The partial differential equation is augmented by periodic boundary conditions along the *x*-direction. In the *y*-direction, we chose no-flux boundary conditions, *n*·∇*µ* = 0, and neutral interactions with the wall, captured byn.. ∇ *c* = 0, where ***n*** is the normal vector at the boundary.

Cluster formation is driven by phase separation, which can be described by the standard free energy 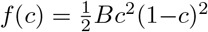 where *f* (*c*) is the free energy density describing local interactions of particles, whereas *κ* penalizes gradients and leads to surface tension effects. For simplicity, we here consider a simple polynomial form, *f* 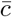with positive energy coefficient *B*. This free energy density exhibits minima at *c*_*−*_ = 0 and *c*_+_ = 1, which correspond to coexisting phases of different density. If the average concentration 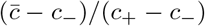in the system is between *c*_*−*_ and *c*_+_, this system will generally exhibit one dense phase, which covers a fraction 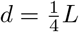of the entire system.

To suppress coarsening, we introduce a region of width *d* at one boundary, where we produce and degrade material to bias the concentration toward *c*_0_. We describe the production of material using the source/sink term

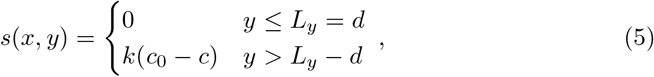

where *k* denotes the reaction rate that determines how fast the system relaxes toward *c*_0_ in the boundary region. For the simulations shown in the main text, we picked *L*_*x*_ = 3*L, L*_*y*_ = *L*, and 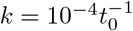.

### Cluster identification

To identify clusters in both simulation and experimental results, we used the DBSCAN clustering method. Properly selecting the DBSCAN parameters—*ϵ* (the maximum distance between points to be considered neighbors) and *N*_*min*_ (the minimum number of points required to form a cluster)—is crucial.

To select the optimal epsilon for the simulation results, we used the elbow technique, which is based on the nearest neighbor algorithm. This method involves plotting the distance to the k-th nearest neighbor for each data point, and identifying the “elbow” point on the graph where the rate of change slows down. This point is typically considered the best choice for *ϵ*. The value of *N*_*min*_ was set individually for each case based on the characteristics of the data.

To ensure accurate parameter selection for imaging data, we defined an error estimation method in which we repeated the same procedure as simulation results to optimize these parameters for multiple cultures (details provided in the supporting information).

## Acknowledgements

We thank Prof. Stefan Klumpp and Prof. Silvio Rizzoli for discussions. We thank Dr. Victor Macarr’on-Palacios for the support with MINFLUX images acquisition. This work was supported by the German Research Foundation (Deutsche Forschungsgemeinschaft, DFG) through grants TE 1172/7-1 [CT], SFB1286 subprojects C01, Z01 [CT], and A07 [ED]. DZ and RR gratefully acknowledge funding from the Max Planck Society and the European Union (ERC, EmulSim, 101044662).

